# MetaCycle: an integrated R package to evaluate periodicity in large scale data

**DOI:** 10.1101/040345

**Authors:** Gang Wu, Ron C. Anafi, Michael E. Hughes, Karl Kornacker, John B. Hogenesch

## Abstract

**Summary:** Detecting periodicity in large scale data remains a challenge. Different algorithms offer strengths and weaknesses in statistical power, sensitivity to outliers, ease of use, and sampling requirements. While efforts have been made to identify best of breed algorithms, relatively little research has gone into integrating these methods in a generalizable method. Here we present MetaCycle, an R package that incorporates ARSER, JTK_CYCLE, and Lomb-Scargle to conveniently evaluate periodicity in time-series data.

**Availability and implementation:** MetaCycle package is available on the CRAN repository (https://cran.r-project.org/web/packages/MetaCycle/index.html) and GitHub (https://github.com/gangwug/MetaCycle).

**Supplementary information:** Supplementary data are available at Bioinformatics online.

## 1 INTRODUCTION

Identifying periodic signals in time-series data is important in studying oscillatory systems, such as the circadian clock or cell cycle. Researchers have devised and evaluated many methods (see a short list in Doherty and Kay, 2010), each with strengths and weaknesses (e.g. Deckard, et al., 2013; Wu, et al., 2014). For example, Fisher’s G Test (Wichert, et al., 2004) and JTK_CYCLE (Hughes, et al., 2010) offer great statistical power to identify rhythmicity, which facilitates correcting for multiple testing in large datasets. On the other hand, some powerful methods, including Fisher’s G Test and ARSER (Yang and Su, 2010), fail without complete, evenly sampled time series data. To avoid such ‘mode failure’ and capture the most value from data, one could choose the best algorithm for each experiment. However, for any single experiment the choice of optimal algorithm is often unclear and may depend on unknown features of the data, such as the shape of the oscillatory patterns or the nature of the noise. Alternatively, a fault-tolerant method that avoids mode failure would have wider applicability, insulating the user from tricky decisions.

Here we present MetaCycle, an N-version programming (NVP) method to explore periodic data. NVP (Avizienis and Chen, 1977) is a concept from the aeronautics industry. Aircraft guidance systems employ multiple independent algorithms and a voting scheme to integrate their results. A method that performs poorly in a particular condition will be outvoted by other methods that better accommodate that condition. NVP has three elements: i) an initial specification that addresses the problem to be solved, data formats, and methods of integrating specified variables from N-version programs, ii) two or more independent algorithmic approaches (N-version software, or NVS) to solve the problem, and iii) an execution environment (N-version executive, or NVX) that runs NVS and provides decision algorithms (Avizienis, 1995; Chen and Avizienis, 1978).

MetaCycle incorporates these concepts into its meta2d mode, which analyzes data from a single time series. Using the same input file, MetaCycle::meta2d implements ARSER (ARS), JTK_CYCLE (JTK), and Lomb-Scargle (LS; Glynn, et al., 2006), from dramatically different disciplines, computer science, statistics, and physics, respectively. After running these algorithms, meta2d integrates their results in a common output file in the (R) environment. We show that MetaCycle::meta2d avoids mode failure while providing robust power in rhythm detection and accurate phase estimations. Further, researchers often want to integrate results from multiple individuals, preserving individual contributions to an overall result. For analyzing data from multiple time series (e.g. several individuals), MetaCycle provides another mode, MetaCycle::meta3d, a convenient and efficient tool to integrate multiple large-scale time series datasets from individual subjects.

## 2 METHODS

Our functional requirement, the initial specification step of NVP, is identifying periodic signals from time-series datasets. At this step, we also specify input data formats, comparison vectors (c-vectors; including p-values, period length, and phase), the decision algorithm, and where it will be applied (cc-points). A detailed description of the decision algorithm is in the vignette. In brief, as default, Fisher’s method (Fisher, 1925) is used to integrate p-values. Arithmetic and circular means are applied in period and phase integration, respectively. The c-vectors do not include amplitude, which are re-calculated with the ordinary least squares method by constructing a general periodic model. Three independent algorithms (ARS, JTK, and LS) were then selected from best of breed methods (Deckard, et al., 2013; Wu, et al., 2014) and developed into NVS. At the third step, we used the R language as the N-version execution environment (NVX). R assures input consistency for each version, local supervision, and implementation of the decision algorithm.

## 3 RESULTS AND DISCUSSION

In model organism experiments, genetically similar or identical animals are typically used, making the identification of each individual sample unimportant. With human research, though, individual subjects are usually recruited, with samples (e.g. blood, skin) collected from the same person over time. Unless samples are pooled, this generates multiple time series datasets. We classify single time-series datasets from one individual or pools as 2D, while multiple time series datasets are designated 3D (Fig. 1). Two functions-*meta2d* and *meta3d* in MetaCycle are designed to analyze time-series datasets in these categories. MetaCycle allows users to select one or more cycling detection algorithm from ARS, JTK and LS to analyze their datasets. Using the same input data format, analyzing different kinds of time-series datasets (evenly or unevenly sampled, with or without replicates or missing values) becomes much easier. Additionally, when analyzing single time-series datasets with two or three algorithms or analyzing multiple time series with the same algorithm, MetaCycle reports integrated results (e.g. p-values).

**Figure 1.**
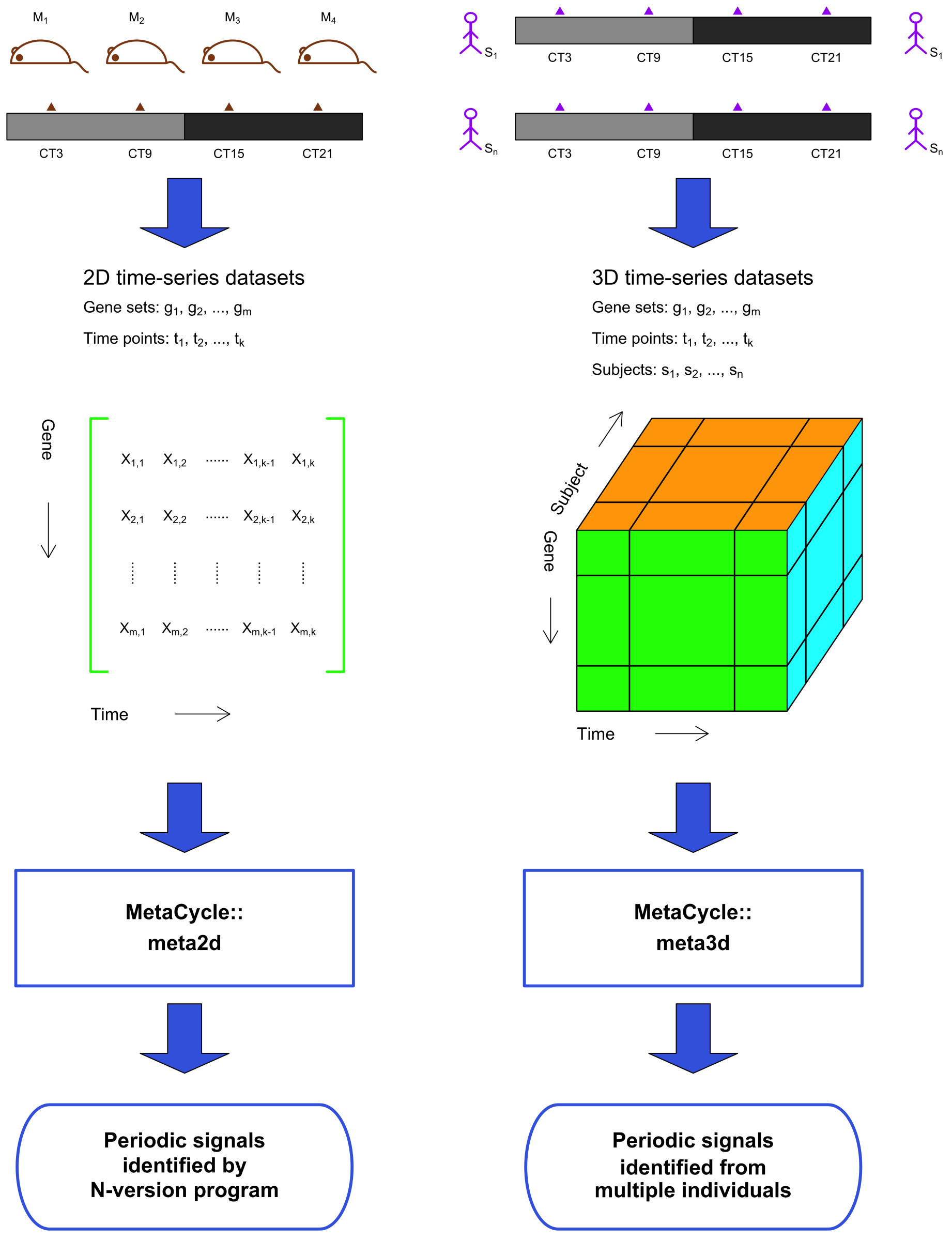
A schematic diagram showing the application of MetaCycle

Using simulated data generated from several different oscillatory reference waves and adding normally distributed noise, we show that NVP generally performs better than any single algorithm in reporting cycling (Fig. S1) or phase values (Fig. S2). In rare cases, NVP may not give better results than the single most suitable method. For example, in analyzing time-series datasets with high temporal resolution (every 1h within 2days), NVP does better than ARS but not as well as LS or JTK (In general, with these conditions, most algorithms do well.). These results indicate that NVP usually gives the best results but will rarely give the worst results. We also applied the NVP method on experimental time-series data from yeast (Orlando, et al., 2008), the NVP method specifically identified 377 clearly periodic genes that were missed in the original analyses (Fig. S3).

With more and different large-scale time-series datasets being produced (e.g. ChIP-seq, proteomics, metabolite profiling), tools for better handling these data are needed. Selecting the best algorithm for each dataset can be tricky, as experimental designs and biological unknowns can interact to produce a myriad of complications. Further, algorithms often require different input formats and run-time parameters and sometimes are implemented in different languages (e.g. in R, Python, or Java). To address these problems, we have developed an NVP method using three popular methods -- ARS, JTK and LS -- in MetaCycle package in R. To facilitate its use and further development, we have made the source code available via Github (https://github.com/gangwug/MetaCycle) and the program available through R repositories (https://cran.r-project.org/web/packages/MetaCycle/index.html).

## ACKNOWLEDGEMENTS

The authors thank Dr. Rendong Yang for his input on applying ARSER, Siqi Liu and Dr. Ben Voight helpful discussions about Fisher’s method for integrating p-values, and Dr. Steve Haase information about his yeast dataset. We also thank members of the Hogenesch lab for valuable discussion and advice on this project.

Funding: This work is supported by the National Institute of Neurological Disorders and Stroke (5R01NS054794-08 to JBH) and the Defense Advanced Research Projects Agency (DARPA-D12AP00025, to John Harer, Duke University).

Conflict of Interest: none declared.

